# H3.3 contributes to chromatin accessibility and transcription factor binding at promoter-proximal regulatory elements

**DOI:** 10.1101/2022.06.30.498282

**Authors:** Amanuel Tafessu, Ryan O’Hara, Sara Martire, Altair L. Dube, Purbita Saha, Laura A. Banaszynski

## Abstract

**Background:** The histone variant H3.3 is enriched at active regulatory elements such as promoters and enhancers in mammalian genomes. These regions are highly accessible, creating an environment that is permissive to transcription factor binding and the recruitment of transcriptional coactivators that establish a unique chromatin post-translational landscape. How H3.3 contributes to the establishment and function of chromatin states at these regions is poorly understood.

**Results:** We performed genomic analyses of features associated with active promoter chromatin in mouse embryonic stem cells (ESCs) and found evidence of subtle yet widespread promoter dysregulation in the absence of H3.3. Loss of H3.3 deposition at promoters reduces chromatin accessibility and transcription factor (TF) footprinting for nearly all TFs expressed in ESCs. H3.3 deletion leads to reduced promoter enrichment of the transcriptional coactivator and histone acetyltransferase, p300. Subsequently, histone H3 acetylation at lysine 27 (H3K27ac) is reduced at promoters in the absence of H3.3, along with reduced enrichment of the bromodomain-containing protein BRD4, an acetyl lysine reader. Despite the observed chromatin dysregulation, H3.3 KO ESCs maintain transcription from ESC-specific genes. However, upon undirected differentiation, H3.3 KO cells retain footprinting of ESC-specific TFs motifs and fail to generate footprints of lineage-specific TF motifs, in line with their diminished capacity to differentiate.

**Conclusions:** H3.3 facilitates DNA accessibility, TF binding, and histone post-translational modification at active promoters. While H3.3 is not required for maintaining transcription in ESCs, it is required for TF binding at new promoters during differentiation.

## Background

In eukaryotic cells, DNA is wrapped around histone proteins to form nucleosomes, the fundamental repeating unit of chromatin [1,2]. While chromatin functions in part to organize a large amount of genomic material within the confines of the nucleus, the natural consequence of this condensation is that regulatory DNA sequences become masked to transcription factors and other proteins that must locate their target sequences for downstream function [3,4]. A subset of specialized transcription factors are able to engage nucleosomal DNA, so-called “pioneer” factors [5]. However, many transcription factors must cooperate with chromatin remodeling factors and the local chromatin environment to engage their target DNA sequences [6]. In addition to specific post-translational modifications, nucleosomes at active regulatory elements are enriched with the histone variants H2A.Z and H3.3 [7]. These nucleosomes are proposed to have unique physical properties that may destabilize the nucleosome core particle [8,9], providing a “window of opportunity” for access to the underlying DNA. Studies of H2A.Z function largely support this view, attributed to both primary sequence differences from replication-coupled H2A and coordinated nucleosome eviction and exchange by dedicated H2A.Z interacting proteins [10–12]. The contribution of H3.3 to chromatin accessibility and transcription factor binding, however, is less clear [13].

H3.3 differs from replication-coupled H3 by only 4-5 amino acids, yet this is sufficient to drive dedicated chaperone association and deposition at specific regions of the genome [7]. H3.3 was first identified as a component of active chromatin [14] and many genome-wide studies have noted its deposition at genic regions such as enhancers, promoters, and gene bodies [15,16]. Regions of H3.3 deposition are sites of dynamic nucleosome turnover [17–21], and several studies have suggested that H3.3 deposition itself may function to destabilize nucleosomes [8,22]. However, other studies have found that H3.3 nucleosomes are structurally and thermodynamically indistinguishable from nucleosomes containing canonical H3 [23,24]. Further, although H3.3 is enriched at active enhancers, previous data show minimal disruption of chromatin accessibility at enhancers in the absence of H3.3, suggesting that H3.3 is correlative with chromatin dynamics rather than causative in this setting [25,26].

Several studies suggest that H3.3 may influence the local chromatin environment by recruiting specific complexes to chromatin [27–29]. For example, H3.3 recruits chromatin remodeling complexes, particularly SWI/SNF and NuRD, whose role in regulating nucleosome dynamics at regulatory elements may influence transcription factor binding [27,30]. In addition, H3.3 has been shown to contribute to the post-translational modification state found at specific regions [7,25,28,29]. For example, recent studies demonstrate that histone H3 lysine 27 acetylation (H3K27ac), a hallmark of active enhancers and promoters thought to occur downstream of transcription factor binding [13,31], is reduced in the absence of H3.3 [25,32,33]. Perhaps surprisingly, H3.3-mediated reduction of enhancer acetylation is not correlated with global reduction of transcription in embryonic stem cells (ESCs) [25,34]. However, a number of studies in mammalian cell lines suggest that H3.3 plays a role in *de novo* transcription [35,36] in response to extracellular stimuli [25,37–41], and H3.3 knockout in animal models results in embryonic lethality or sterility [42–44]. Together, these observations suggest that H3.3 may be functionally important to initiate new transcription programs.

In this study, we performed genomic analyses to determine the effect of H3.3 deposition on regulatory element architecture and downstream transcription in mouse embryonic stem cells (ESCs). We find that promoter-proximal regulatory elements become less accessible in the absence of H3.3. Reduced accessibility is accompanied by reduced transcription factor footprinting, attenuation of chromatin states thought to be downstream of transcription factor binding, and decreased RNA polymerase II engagement at affected promoters. While ESCs appear quite tolerant to these changes in genome regulation, they are unable to respond to cellular differentiation cues and show perdurance of the regulatory landscape associated with pluripotency. Thus, we propose that H3.3 is a necessary upstream component of transcriptional activation that becomes dispensable for the maintenance of established gene regulatory networks.

## Results

### H3.3 increases DNA accessibility at promoters

Given the dynamic transcription-associated turnover of H3.3 at regulatory elements [20], we wanted to test whether H3.3 is required for accessibility at transcriptionally active regions. We first performed H3.3 ChIP-seq and ATAC-seq on WT mouse embryonic stem cells (ESCs) [25]. All ATAC-seq studies were performed in technical duplicate at a minimum sequence depth of 40 million reads for each data set. In agreement with previous studies, we find that H3.3 enrichment at active regulatory elements is correlated with accessibility (Fig. 1A, Fig. S1A). Accessibility at active promoters (defined as >20 baseMean across WT and H3.3 KO RNA-seq [25]) showed higher correlation with H3.3 deposition compared to active promoter-distal regulatory elements (defined as regions +/− 3 kb from a promoter and containing both ATAC-seq and H3K27ac ChIP-seq enrichment). We next asked whether regulatory element accessibility is dependent upon H3.3 by comparing ATAC-seq data from WT and H3.3 KO ESCs [25,29]. In line with higher correlation between accessibility and H3.3 deposition at promoters, we observe a slight but significant decrease in chromatin accessibility at promoters but not distal regulatory elements in the absence of H3.3 (Fig. 1B,C and Fig. S1B). Decreased promoter accessibility is apparent both at the level of individual promoters and genome-wide (Fig. 1B-C). We next wanted to determine whether reduced promoter accessibility in H3.3 KO ESCs is related to the level of H3.3 enrichment at that promoter in WT ESCs. Globally, we find that reduced ATAC-seq signal at promoters in H3.3 KO ESCs is correlated with H3.3 enrichment in WT ESCs (Fig 1D). Further, by binning promoters into quartiles based on H3.3 enrichment, we observed that promoters with higher H3.3 enrichment indeed showed more pronounced and significant loss of accessibility by ATAC-seq in H3.3 KO ESCs (Fig. 1E).

**Figure 1.**
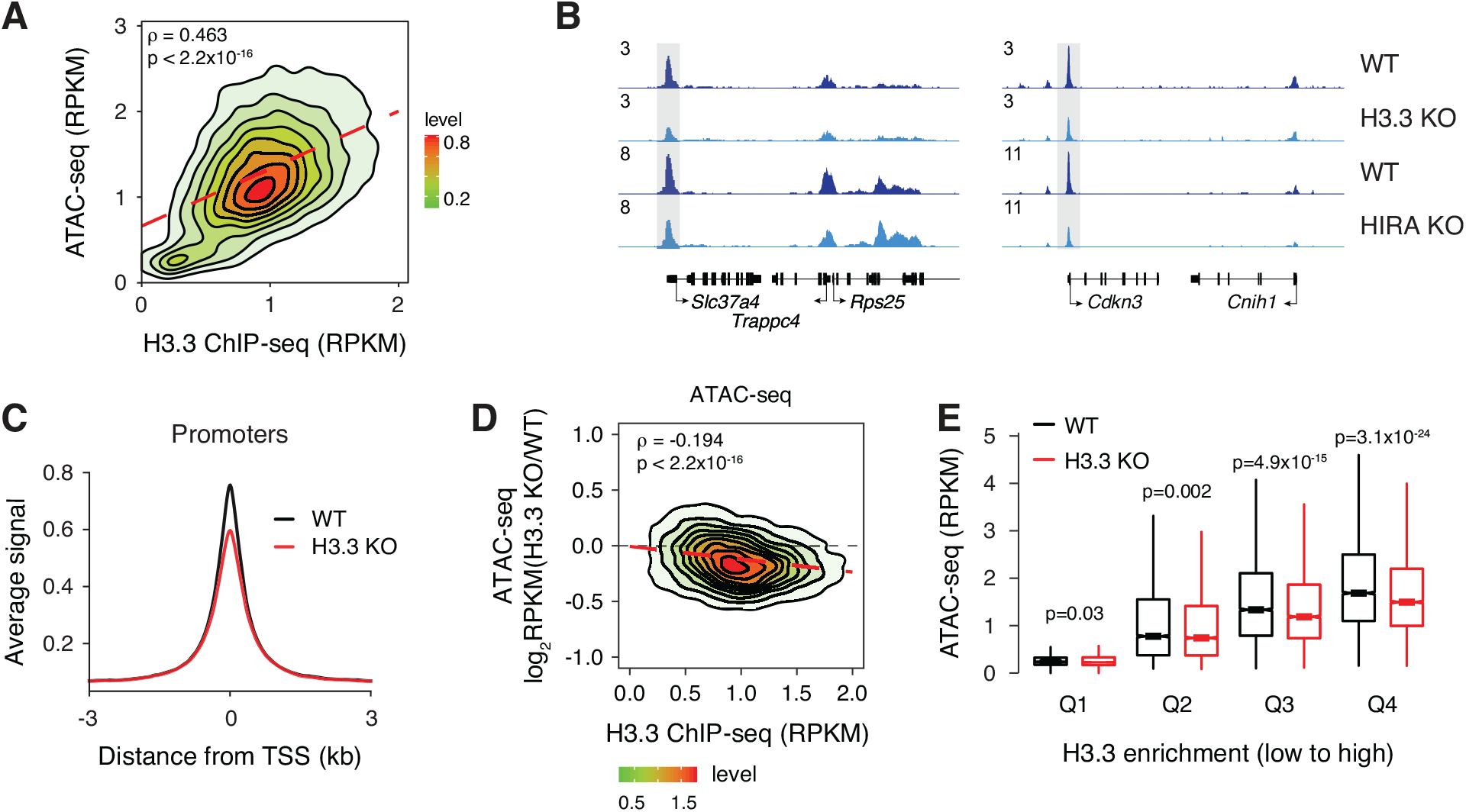
Loss of H3.3 reduces chromatin accessibility at promoters. **A** Correlation plot between ATAC-seq and H3.3 ChIP-seq at promoters in ESCs. **B** Genome browser representations of ATAC-seq in WT and H3.3 KO ESCs and WT and HIRA KO ESCs. The y-axis represents read density in reads per kilobase per million mapped reads (RPKM). **C** ATAC-seq average profiles at promoters in WT and H3.3 KO ESCs. **D** Correlation plot between differential ATAC-seq signal in H3.3 KO compared to WT ESCs and H3.3 enrichment at promoters in WT ESCs. **E** Boxplot showing ATAC-seq signal at promoters binned by H3.3 enrichment in WT and H3.3 KO ESCs. The bottom and top of the boxes correspond to the 25th and 75th percentiles, and the internal band is the 50th percentile (median). The plot whiskers correspond to 1.5x interquartile range and outliers are excluded. P-values determined by Wilcoxon rank sum two-side test.

H3.3 deposition occurs through two distinct chaperone complexes. The HIRA complex is responsible for the majority of H3.3 deposition at promoters, gene bodies, and enhancers, whereas the ATRX–DAXX complex deposits H3.3 at repetitive regions such as telomeres and interstitial heterochromatin [7,15]. We therefore predicted that loss of HIRA, but not ATRX or DAXX, would result in similar effects on promoter accessibility as observed upon loss of H3.3. In agreement, we observed a similar reduction in promoter accessibility in HIRA KO ESCs (Fig. 1B, Fig. S1C-D). By contrast, ATRX KO did not alter regulatory element accessibility and DAXX KO interestingly resulted in a slight increase in chromatin accessibility at both enhancers and promoters (Fig. S1E-I). Genome-wide, we identified 335 regions of differential accessibility at promoters in H3.3 KO compared to WT ESCs (p < 0.05), with 79% (265/335) of these regions becoming less accessible in H3.3 KO ESCs (Fig. S2A). These “H3.3-dependent” regions were also less accessible in HIRA KO ESCs but not in DAXX KO or ATRX KO ESCs (Fig. S2B). In addition to reduced accessibility, we find that loss of either H3.3 or HIRA, but not ATRX or DAXX, disrupts nucleosome footprinting and positioning at promoters genome-wide as assessed by NucleoATAC [45] (Fig. S3). Overall, these data suggest that HIRA-dependent deposition of H3.3 at promoters has a role in maintaining chromatin accessibility and nucleosome organization at these regions.

### H3.3 facilitates transcription factor binding at promoters

Given that chromatin accessibility is a hallmark of regulatory element transcription factor binding, we hypothesized that the reduced promoter accessibility observed in H3.3 KO and HIRA KO ESCs would be accompanied by reduced TF binding. ATAC-seq data measures chromatin accessibility but also contains regions of depleted signal within open chromatin that are protected from transposition by TF binding, referred to as TF footprints. We used a recently developed tool called TOBIAS [46] to perform comparative footprinting analysis of 395 expressed TFs with known consensus motifs in WT and H3.3 KO ESCs. This analysis revealed that nearly all expressed TFs show reduced binding to their motifs in the promoters of H3.3 KO ESCs, including key members of the transcription factor network that controls pluripotency, POU5F1 (e.g., OCT4, SOX2, and Nanog [47]) (Fig. 2A, Table S1). Analysis of ATAC-seq data from WT and H3.3 KO ESCs centered on motifs for selected pluripotency TFs showed clearly reduced accessibility upon loss of H3.3 (Fig. 2B). While all TFs show reduced binding scores in the absence of H3.3, those families most affected based on the magnitude and significance of dysregulation bind to motifs containing high GC content (Fig. 2C, Fig. S2C), perhaps reflective of their presence at promoters. In line with our accessibility analysis, HIRA KO ESCs showed a similar reduction of TF binding while DAXX KO and ATRX KO ESCs did not (Fig. S4, Table S1). As expected, the most highly dysregulated TF families in H3.3 KO and HIRA KO ESCs overlap significantly (Fig. 2D).

**Figure 2.**
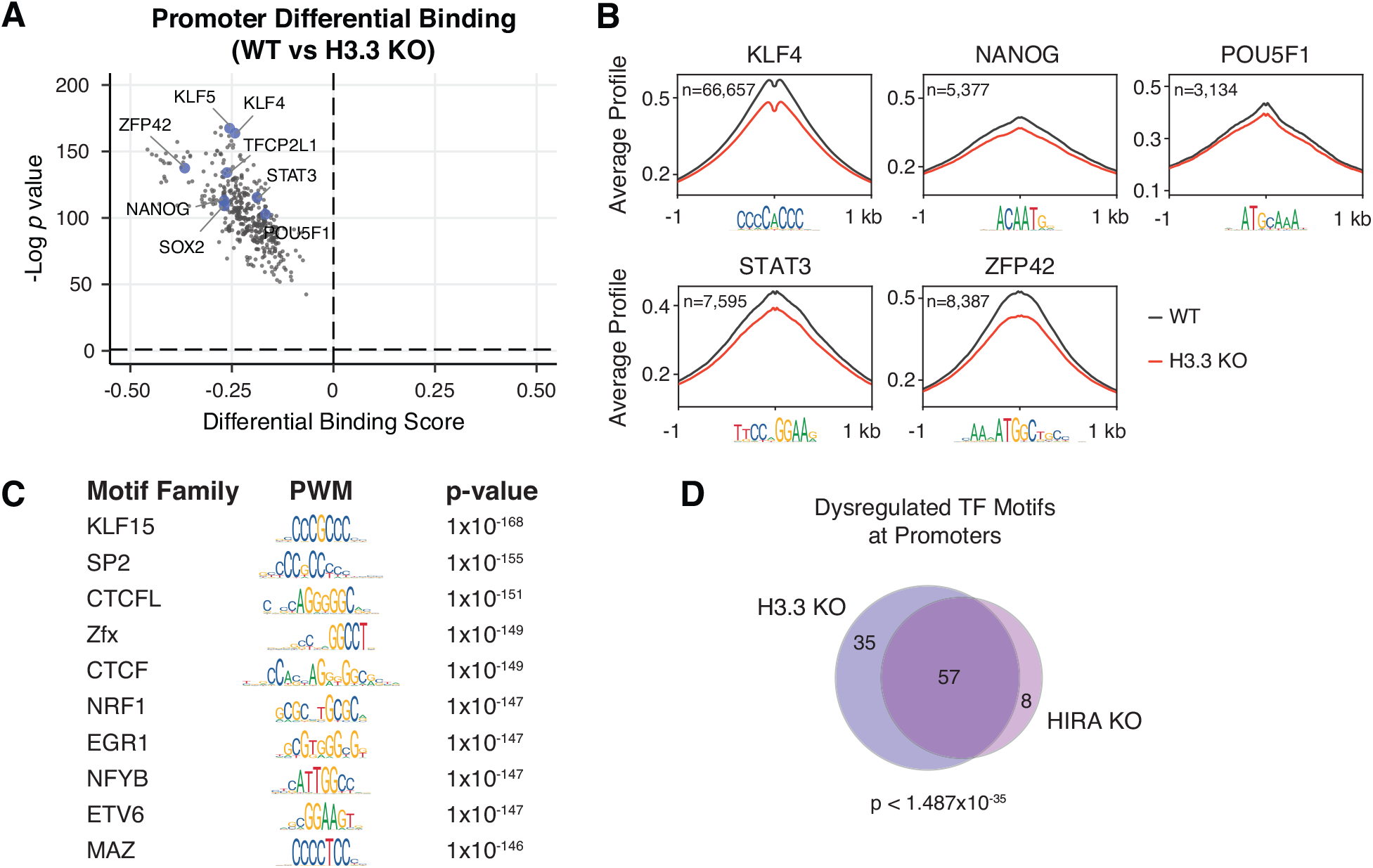
Loss of H3.3 reduces TF footprinting at promoters. **A** Pairwise comparison of TF activity at promoters between WT and H3.3 KO ESCs. The volcano plot shows differential binding activity against the -log10(p value) for all investigated TF motifs. Each TF is represented by a single circle (n = 395). TF motifs enriched in WT ESCs have negative differential binding scores and TF motifs enriched in H3.3 KO ESCs have positive differential binding scores. Motifs for a subset of pluripotency-associated TFs are highlighted in blue. **B** ATAC-seq average profiles at representative TF motifs at promoters in WT and H3.3 KO ESCs. Data are centered on the motif and the number of motifs profiled are indicated. **C** Illustration of motifs of the most dysregulated TF families at promoters in H3.3 KO ESCs classified based on Manhattan scores in the top 10% across all comparisons (i.e., WT vs H3.3 KO, HIRA KO, ATRX KO, or DAXX KO). **D** Venn diagram representing TF motifs commonly dysregulated at promoters in H3.3 KO and HIRA KO ESCs based on Manhattan score as described above.

Loss of TF binding in H3.3 and HIRA KO ESCs could be caused either by altered binding ability or by reduced TF expression. When we compared published RNA-seq data for WT, H3.3 KO and HIRA KO ESCs [25], we did not observe a global loss of TF expression in either H3.3 KO or HIRA KO ESCs (Fig. S5A,B; Table S2). Further, we found no correlation between changes in TF expression and differential TF binding scores due to loss of H3.3 or HIRA (Fig S5C,D). Finally, we do not observe changes in expression at the protein level for a subset of assessed pluripotency TFs in H3.3 KO compared to WT ESCs (Fig. S5E). Together, our results suggest that loss of H3.3 deposition does not affect TF levels but rather affects the ability of TFs to bind to promoters.

### Dysregulation of promoter chromatin landscape in the absence of H3.3

Promoters of active genes contain distinct chromatin post-translational modifications. These regions are enriched with H3K27ac and H3K4me3, deposited by the CBP/p300 acetyltransferases and the MLL1/2 methyltransferases, respectively [48,49]. Recruitment of these enzymes has been shown to be downstream of TF binding [13,50–53]. CBP/p300- mediated histone acetylation in turn acts as a scaffold to recruit effector proteins such as the bromodomain and extra-terminal domain (BET) family protein BRD4, which is involved in transcription elongation [54]. Given the reduction in both accessibility and TF binding observed at promoters lacking H3.3, we hypothesized that subsequent steps in establishing the promoter chromatin landscape may be dysregulated.

To explore the relationship between H3.3 deposition and chromatin signatures at promoters, we both reanalyzed existing data sets [25] and performed additional chromatin immunoprecipitation followed by sequencing (ChIP-seq) of several modifications and chromatin-associated proteins in WT and H3.3 KO ESCs. Genome-wide, we observed only a subtle decrease in H3K4me3 at active promoters (Fig. 3A-B, Fig. S6A,E), suggesting that this modification has little reliance on H3.3 for its installation into chromatin. In contrast, the molecular machinery associated with histone acetylation shows greater dependence on H3.3 at promoters. We find that loss of H3.3 leads to reduced enrichment of p300, H3K27ac, and BRD4 at active promoters genome-wide (Fig. 3A,C-E, Fig. S6B-D). Reduced enrichment of each factor in H3.3 KO ESCs was correlated with the level of H3.3 enrichment present at WT promoters (Fig. S6F-H). The effect of H3.3 loss on p300, H3K27ac, and BRD4 was not restricted to promoters containing specific TF motifs, but rather appeared more global, with even regions bound by lineage-drivers such as Oct4, Sox2, and Nanog showing reduced enrichment (Fig. 3F, Fig. S6I). Reduced recruitment of p300 and BRD4 at promoters cannot be attributed to reduced expression of these genes in H3.3 KO ESCs (Fig. S5), suggesting that the presence of H3.3 itself plays a role in regulating histone acetylation at promoters.

**Figure 3.**
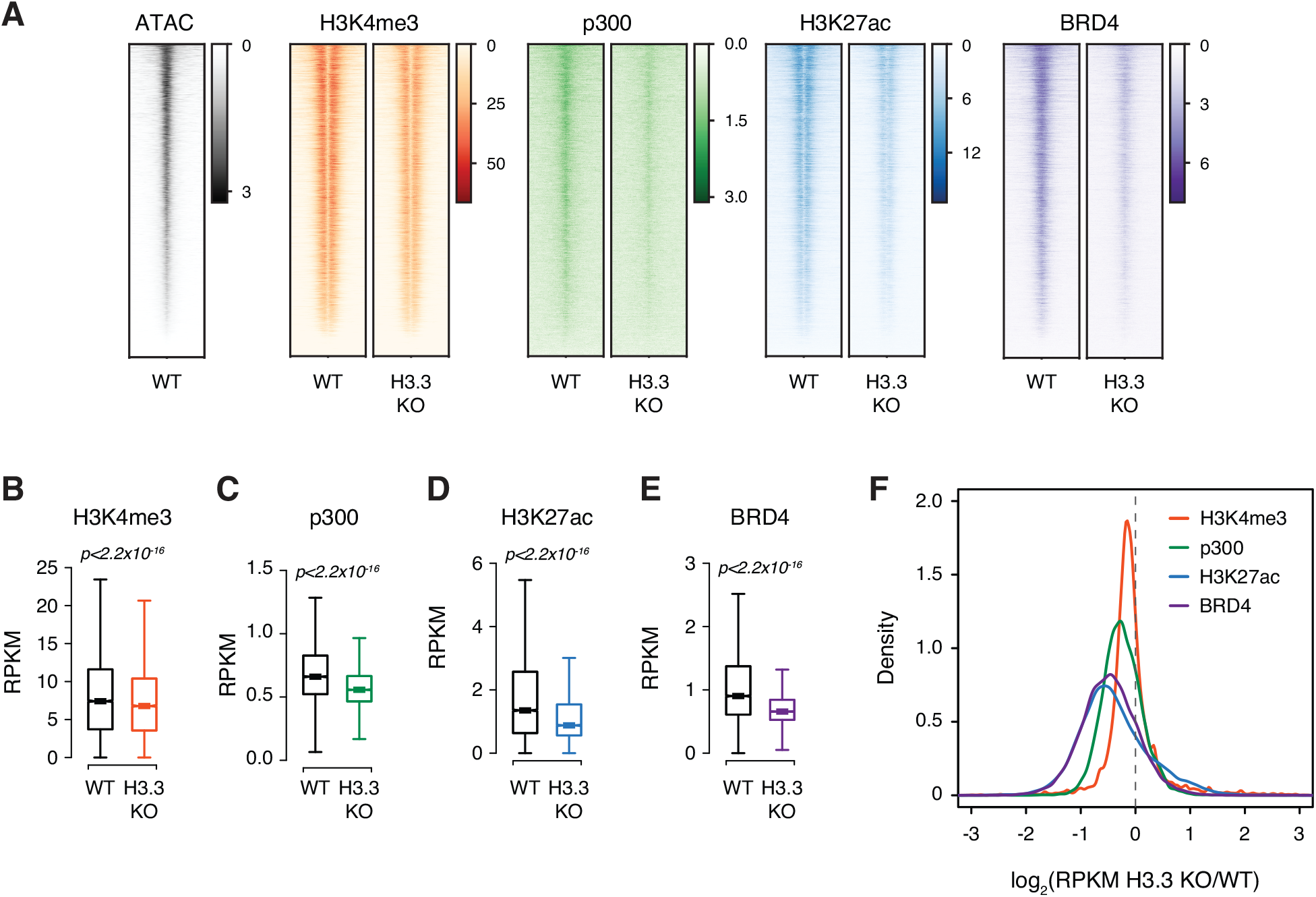
Chromatin landscape is dysregulated with H3.3 loss. **A** Heatmaps of ATAC-seq in WT and H3K4me3, p300, H3K27ac, and BRD4 enrichment at promoters in WT and H3.3 KO ESCs. 3 kb around the center of promoters are displayed for each analysis. Each row represents a single active promoter (n = 12,903). **B-E** Boxplots showing **(B)** H3K4me3, **(C)** p300, **(D)** H3K27ac, and **(E)** BRD4 enrichment at active promoters in WT and H3.3 KO ESCs (n = 12,903). The bottom and top of the boxes correspond to the 25th and 75th percentiles, and the internal band is the 50th percentile (median). The plot whiskers correspond to 1.5x interquartile range and outliers are excluded. P-values determined by Wilcoxon rank sum two-side test. **F** Ratio (log2) of H3K4me3, p300, H3K27ac, and BRD4 enrichment at promoters in WT and H3.3 KO ESCs. x axis values <0 indicate reduced enrichment in the absence of H3.3.

Finally, since H3.3 deposition at promoters is facilitated by the HIRA complex, we again expect that loss of HIRA, but not ATRX or DAXX, will phenocopy the effects of loss of H3.3 at promoters. As proof-of-principle, we reanalyzed existing H3K27ac ChIP-seq data sets obtained from WT, HIRA, ATRX, or DAXX KO ESCs [25]. In agreement with our ATAC-seq results, we find that only loss of HIRA resulted in reduced H3K27ac at active promoters compared to WT ESCs (Fig. S6J). In contrast, loss of ATRX had no effect on promoter H3K27ac enrichment while loss of DAXX resulted in a slight increase in H3K27ac at promoters (Fig. S6K,L). This effect was clear when directly comparing the ratio of H3K27ac enrichment at individual promoters in WT and H3.3 chaperone KO ESCs (Fig. S6M). Overall, our data demonstrate that HIRA-dependent H3.3 deposition positively influences p300 binding, H3K27 acetylation, and recruitment of downstream effectors such as BRD4 at promoters.

### Reduced promoter RNA polymerase II engagement in H3.3 KO cells

Our results demonstrate that promoters lacking H3.3 show reduced enrichment of histone modifications and cofactors characteristic of transcriptional activity (Fig 3), suggesting that this activity may be reduced in the absence of H3.3. To test whether loss of these marks was associated with reduced active RNA polymerase II (RNAPII) engagement, we performed global run-on sequencing (GRO-seq) from WT and H3.3 KO ESCs. This technique relies on the strong interaction between transcriptionally engaged RNAPII and DNA to produce nascent transcripts in vitro that are then sequenced and mapped to determine sites of active RNAPII within the genome [55]. In agreement with dysregulation of the chromatin landscape at active promoters, we find that active RNAPII engagement was reduced at the TSS of expressed genes in H3.3 KO compared to WT ESCs (Fig 4A). Interestingly, the change in GRO-seq signal at individual promoter-proximal TF motifs in H3.3 KO compared to WT ESCs was correlated with reduced footprinting of that TF in H3.3 KO ESCs, suggesting an association between the magnitude of TF dysregulation and the reduction of RNAPII engagement (Fig 4B).

**Figure 4.**
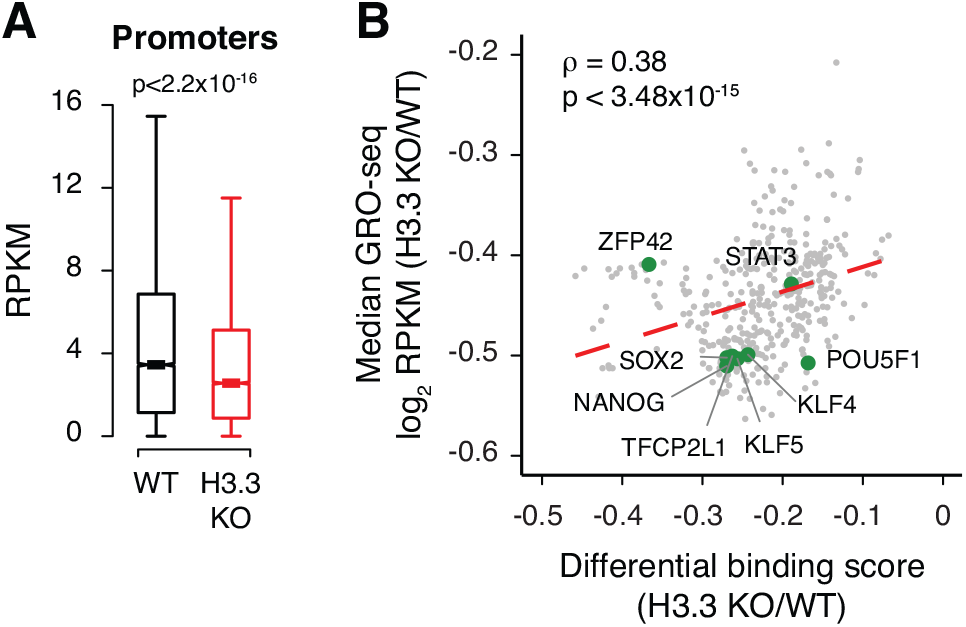
H3.3 facilitates active pol II engagement at promoters. **A** Box plot showing GRO-seq signal at promoters of expressed genes (TSS - 30 bp to TSS + 250 bp) in WT and H3.3 KO ESCs. The bottom and top of the boxes correspond to the 25th and 75th percentiles, and the internal band is the 50th percentile (median). The plot whiskers correspond to 1.5x interquartile range and outliers are excluded. P-values determined by Wilcoxon rank sum two-side test. **B** Scatterplot showing differential binding scores of investigated TF motifs and median ratio (log2) of GRO-seq signal at promoters containing each motif. Each TF is represented by a single dot (n = 395). Representative TF motifs are labelled in green. Dashed red line represents a linear fit.

### H3.3 is required for TF activity during differentiation

Surprisingly, loss of RNAPII engagement observed in H3.3 KO ESCs is not associated with a global reduction in steady-state transcription, as measured by RNA-seq [25]. Although we do observe differentially expressed genes by RNA-seq, the core ESC transcriptional program remains mostly unperturbed. Of the 335 genes that constitute the “core ESC-like gene module” [56], only 8 genes show >2-fold reduced expression in H3.3 KO ESCs (Fig S7A). Similarly, expression of the PluriNet protein-protein network characteristic of pluripotent cells [57] is maintained in the absence of H3.3 (Fig S7B). Thus, although RNAPII engagement is reduced in the absence of H3.3, the remaining RNAPII appears sufficient for maintaining the ESC transcriptome.

While H3.3 KO ESCs maintain their ability to self-renew, we previously reported that H3.3 is required for undirected differentiation of ESCs into embryoid bodies (EBs) [25]. This process requires both the decommissioning of the existing transcription program as well as the establishment of new gene regulatory networks driven by lineage-specific TFs [58]. H3.3 KO ESCs show a defect in EB formation [25] accompanied by failure to down-regulate ESC-specific genes (Fig. S7C,D). Given the reduced TF binding scores observed in H3.3 KO ESCs, we hypothesized that H3.3 may be involved in the establishment of new lineage-specific TF binding during EB formation.

The complex mixture of cell types present in EBs poses a challenge for TF footprint analysis. To test whether our approach can distinguish between ESC and EB-specific TF footprinting, we generated ATAC-seq data sets from WT ESCs and WT cells differentiated for four days into EBs and performed footprint analysis using TOBIAS [46]. Using our previously published EB and ESC RNA-seq data [25], we compiled a list of 458 TFs which were expressed in either EB or ESCs, including 63 EB-specific and 16 ESC-specific TFs. Since we were specifically interested in the effect of H3.3 loss on the decommissioning of ESC genes and on the activation of EB transcriptional programs, we restricted our analysis to promoters which were expressed specifically in either cell state, resulting in 8,389 ATAC-seq peaks that were unique to either EBs or ESCs. As expected, TF motifs characteristic of pluripotent cells (e.g., KLF4, NANOG, and SOX2) were more bound in ESCs, whereas lineage-specific transcription factors (e.g., FOXA2, GATA2, HAND2) had greater footprinting in EBs (Fig 5A, Table S3). This result gave us confidence that we could distinguish between EB and ESC-specific footprinting.

**Figure 5.**
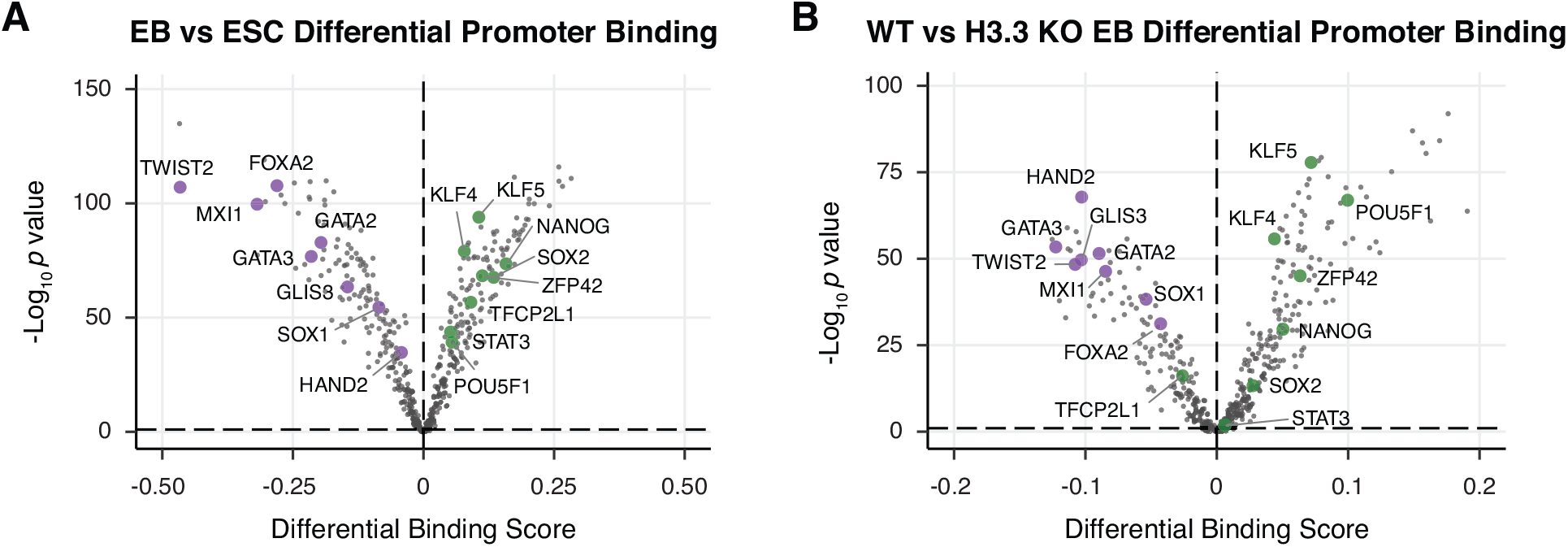
H3.3 supports TF binding during differentiation. **A** Pairwise comparison of TF activity at promoters between WT ESCs and EBs. TF motifs enriched in EBs have negative differential binding scores and TF motifs enriched in ESCs have positive differential binding scores. **B** Pairwise comparison of TF activity at promoters between WT and H3.3 KO EBs. TF motifs enriched in WT EBs have negative differential binding scores and TF motifs enriched in H3.3 KO EBs have positive differential binding scores. For both panels, the volcano plot shows differential binding activity against the -log10(p value) for all investigated TF motifs. Each TF is represented by a single circle (n = 458). Representative differentiation-specific TFs are labeled in purple and representative pluripotency-specific TFs are labeled in green.

To determine whether loss of H3.3 was associated with failure to establish new patterns of TF binding during differentiation, we performed ATAC-seq on WT and H3.3 KO EBs and compared TF footprinting at cell state-specific ATAC-seq peaks. In agreement with the failure of H3.3 KO cells to properly form EBs, our analyses revealed that pluripotency-maintaining TFs tended to remain bound to their motifs (e.g., POU5F1, NANOG, and ZFP42) in H3.3 KO EBs, suggesting an inability to decommission ESC-specific binding patterns. Likewise, lineage-specific TFs (e.g., HAND2, GATA3, and TWIST2) show greater binding in WT EBs, suggesting a defect in initiating differentiation-specific binding events in H3.3 KO cells (Fig 5B, Table S3). Taken together, our analyses suggest that while H3.3-mediated TF binding may not be required to maintain gene regulatory networks in ESCs, H3.3 deposition is essential for establishing new TF binding patterns during differentiation.

## Discussion

The specific contribution of H3.3 to the establishment and maintenance of transcriptionally permissive chromatin remains an open question. Our genomic studies have revealed a relationship between H3.3 deposition and accessibility of active promoters in ESCs. Our data suggest a role for H3.3 in facilitating TF binding to these regions, as well as maintenance of histone modifications and cofactors associated with transcriptionally active genes, including RNAPII engagement. H3.3 KO ESCs are able to maintain transcription despite global, albeit modest, dysregulation of chromatin architecture at promoters. In contrast, loss of H3.3 has dramatic consequences on EB differentiation which coincides with widespread failure to engage lineage-specific TF motifs. Taken together, our data suggests distinct roles for H3.3 in the maintenance of and initiation of transcription.

While the features of active regulatory elements are well-characterized, the order of events that generate transcriptionally permissive chromatin landscapes remains poorly understood. The binding of pioneer TFs to nucleosomal DNA is thought to initiate the assembly of chromatin-modifying protein complexes at promoters [5,25]. However, even pioneer TFs have been shown to be sensitive to nucleosome composition and reliant on specific chromatin remodeling complexes. For instance, the recruitment of OCT4, a model pioneer TF [59,60], is facilitated both by H2A.Z and BAF complex recruitment in ESCs [11,61]. Further, the BAF complex has been shown to facilitate reprogramming of somatic cells to induced pluripotent stem cells (iPSCs) by enhancing OCT4 binding to target sequences [62]. In our assessment, both pioneer factors as well as TFs that are dependent on chromatin remodeling complexes [63] were affected by loss of H3.3 in ESCs, suggesting that this replacement variant is broadly required for optimal TF binding to DNA. Interestingly, a previous study reported that depletion of H3.3 early in reprogramming facilitates the repression of somatic genes; however, H3.3 deposition at later time points was required to initiate ESC-like transcription [64]. These observations are in line with our own and suggest a dual role for H3.3 in both safeguarding cellular identity but also facilitating new transcription during cell fate transitions.

Previous studies have shown that HIRA-dependent H3.3 deposition at promoters proceeds via a gap-filling mechanism to protect the transiently naked DNA that is exposed in the wake of RNAPII transcription [17,21]. One interpretation of our results is that H3.3 itself is necessary for nucleosome displacement to occur. Reduced access to underlying DNA in the absence of H3.3 then results in reduced recruitment of TFs to their target motifs, setting off a cascade of events resulting in reduced p300 recruitment with subsequent downregulation of H3K27ac at promoters. While this seems plausible at promoters, it is important to note that phosphorylation of a unique serine on the H3.3 tail (substituted by an alanine in replication-coupled H3) has been shown to stimulate p300 activity and H3K27ac at both enhancers and promoters [25,33]. In contrast to what we see at promoters, loss of H3K27ac at enhancers occurs without any appreciable decrease in p300 recruitment or change in chromatin accessibility [25], suggesting that distal regulatory elements are subject to distinct mechanisms of control in ESCs.

Recent studies have shown that loss of histone post-translational modifications long associated with active enhancers and promoters has little effect on ongoing transcription [25,34,65,66], with several studies suggesting that histone-modifying enzymes play non-catalytic roles in transcription [67–70]. In line with these findings, we observed that H3.3 KO ESCs were largely able to maintain their transcription program and cell identity despite reduced H3K27ac enrichment at promoters. However, we and others have previously shown a requirement for H3.3 during differentiation [25,32]. In the current study, we showed that loss of H3.3 results in reduced footprinting of lineage-specific TFs and a failure to disengage master ESC regulators during differentiation. Interestingly, proper ESC differentiation requires enhancer decommissioning by the H3K4/K9 demethylase LSD1 [71], a component of the NuRD complex which has been shown to be recruited by H3.3 [27,71]. Thus, it is possible that the persistence of ESC-specific TF networks we observed in H3.3 KO EBs is due to a failure to decommission active enhancers in addition to an inability to recruit TFs to lineage-specific regulatory elements.

Interestingly, H3.3 has previously been shown to facilitate the recruitment of both BAF and NuRD complexes in a manner that requires the H3.3 K4 residue [27,30]. Given the widespread reliance of TFs on chromatin remodeling, it is possible that this function of H3.3 underlies the extensive TF dysregulation we observe in H3.3 KO ESCs. It remains to be seen whether mutation of specific H3.3 residues or deletion of specific chromatin remodelers can recapitulate loss of chromatin accessibility and TF binding that we observe in H3.3 KO ESCs. Future studies investigating the activities of specific remodelers in the absence or mutation of H3.3 will shed light on how H3.3 influences chromatin dynamics and transcriptional regulation.

## Conclusions

In this study, we investigated the contributions of histone H3.3 to chromatin states at promoters. Using genomic analyses, we find that H3.3 promotes accessibility, TF binding, and the enrichment of transcriptional coactivators p300 and BRD4 at active promoters. Active RNAPII and histone acetylation associated with active promoters are also depleted in the absence of H3.3, with seemingly no global effect on steady-state transcription. However, in agreement with previous reports, we find that H3.3 is important for gene regulation during differentiation. Specifically, H3.3 is required for the rewiring of TF networks observed during lineage commitment. Our findings build on previous work linking H3.3 deposition to gene activation and identify a role for H3.3 in maintaining transcriptionally permissive chromatin. Given that H3.3 mutations have been identified in pediatric cancers and congenital neurologic disorders, our studies on normal H3.3 function have important implications towards understanding how dysregulation of this histone variant influences human disease [72–74].

## Methods

### ESC culture

ESCs were maintained under standard conditions on gelatin-coated plates at 37 ℃ and 5% CO_2_, in medium containing Knockout DMEM (Thermo Fisher) supplemented with NEAA, GlutaMAX, penicillin/streptomycin (Thermo Fisher), 10% ESC-screened fetal bovine serum (Hyclone), 0.1 mM 2-mercaptoethanol (Fisher) and leukemia-inhibitory factor (LIF). Generation of H3.3 KO, ATRX KO, DAXX KO, and HIRA KO ESCs has been described previously [25,29,75]. ESCs were routinely screened for mycoplasma. For EB formation, ESCs were diluted to 10^4^ cells/ml in EB differentiation media (DMEM, 15% FBS, 1x MEM-NEAA, 1x Pen/Strep, 50 μM β-mercaptoethanol) and 30 μl drops were placed on the lid of a 150 mm dish. The lid was inverted and placed over a dish containing 10–15 ml of PBS. The hanging drops were cultured for 3 days at 37 ℃ and 5% CO_2_. The hanging drops were then washed from the lids with EB differentiation media and cultured in 100 mm dishes on an orbital shaker at 50 rpm for an additional day.

### Antibodies

Brd4 (A301-985A50, Bethyl), H3 general (ab1791, Abcam, Lot # GR177884-2), H3.3 (09-838, Millipore, Lot # 2578126), H3K4me3 (39159, Active Motif), Spike-In antibody (61686, Active Motif, Lot# 00419007), OCT4 (sc-5279, Santa Cruz), NANOG (ab70482, Abcam), KLF4 (ab34814, Abcam), anti-mouse IgG-HRP (NA93V, GE, Lot # 9773218), anti-rabbit IgG-HRP (170-6515, Biorad, Lot # 350003248).

### Chromatin Immunoprecipitation (ChIP)

#### Native ChIP

Cells were trypsinized, washed and lysed (50 mM TrisHCl pH 7.4, 1 mM CaCl_2_, 0.2% Triton X-100, 10 mM NaButyrate, and protease inhibitor cocktail (Roche)) with micrococcal nuclease (Worthington) for 5 min at 37 °C to recover mono- to tri-nucleosomes. Nuclei were lysed by brief sonication and dialyzed twice into RIPA buffer (10 mM Tris pH 7.6, 1 mM EDTA, 0.1% SDS, 0.1% Na-Deoxycholate, 1% Triton X-100) for 1 hr at 4 °C. Soluble material was combined with 50 ng spike-in chromatin (Active Motif 53083) and 5% was reserved as input DNA. 5 μg of antibody and 2 µg of spike-in antibody (Active Motif 61686) were bound to 50 μl protein A or protein G Dynabeads (Invitrogen) and incubated with soluble chromatin overnight at 4 °C. Magnetic beads were washed as follows: 3x RIPA buffer, 2x RIPA buffer + 300 mM NaCl, 2x LiCl buffer (250 mM LiCl, 0.5% NP-40, 0.5% NaDeoxycholate), 1x TE + 50 mM NaCl. Chromatin was eluted and treated with RNaseA and Proteinase K. ChIP DNA was purified using QIAquick PCR Purification Kit (Qiagen).

#### Crosslink ChIP

WT and H3.3 KO ESCs were harvested and crosslinked with 1% formaldehyde in PBS for 10 min at room temperature. Cross-linking was quenched with 125 mM glycine. Cells were lysed (50 mM HEPES, pH 7.5, 140 mM NaCl, 1 mM EDTA, 10% glycerol, 0.5% NP-40, 0.25% Triton X-100) and nuclei were resuspended in ChIP buffer (10 mM Tris, pH 8, 100 mM NaCl, 1 mM EDTA, 0.5 mM EGTA, 0.1% sodium deoxycholate, 0.5% N-lauroylsarcosine). Chromatin was sonicated to an average size of 0.3–1 kb using a Covaris M220 Focused-ultrasonicator. 50 ng spike-in chromatin (Active Motif 53083) was added to the soluble fraction and incubated with 50 μl Protein A Dynabeads (Invitrogen) bound to 5 μg of BRD4 antibody (Bethyl A301-985A50) and 2 µg spike-in antibody (Active Motif 61686) overnight at 4 °C. Dynabeads were washed once with each of the following: low salt wash buffer (10 mM Tris HCl, pH 8, 2 mM EDTA, 0.1% SDS, 1% Triton X-100, 150 mM NaCl), high salt wash buffer (10 mM Tris HCl, pH 8, 2 mM EDTA, 0.1% SDS, 1% Triton X-100, 500 mM NaCl), LiCl wash buffer (10 mM Tris HCl, pH 8, 1 mM EDTA, 1% NP-40, 1% Na-deoxycholate, 250 mM LiCl), and a final wash with TE + 50 mM NaCl. Chromatin was eluted, incubated overnight at 65℃, treated with RNase A and proteinase K, and DNA was purified using QIAquick PCR Purification Kit (Qiagen).

#### ChIP-seq Library Preparation

ChIP-seq libraries were prepared from 5-10 ng ChIP DNA following the Illumina TruSeq protocol. The quality of the libraries was assessed using a D1000 ScreenTape on a 2200 TapeStation (Agilent) and quantified using a Qubit dsDNA HS Assay Kit (Thermo Fisher). Libraries with unique adaptor barcodes were multiplexed and sequenced on an Illumina NextSeq 500 (paired-end, 33 base pair reads). Typical sequencing depth was at least 20 million reads per sample.

#### ChIP-seq analysis

Quality of ChIP-seq datasets was assessed using the FastQC tool. ChIP-seq raw reads were aligned separately to the mouse reference genome (mm10) and the spike-in drosophila reference genome (dm3) using BWA [76]. Only one alignment is reported for each read (either the single best alignment or, if more than one equivalent best alignment was found, one of those matches selected randomly). Duplicate reads were filtered using Picard. Uniquely mapped drosophila reads were counted in the sample containing the least number of drosophila mapped reads and used to generate a normalization factor for random downsampling. Reads were converted into bigWig files using BEDTools [76,77] for visualization in Integrative Genomics Viewer [78]. Peak calling was performed with MACS2 software [79] using a p value cutoff of 0.01. Heatmaps and average profiles were generated using deepTools. Box plots and density plots representing ChIP-seq read densities and fold-changes in read densities, respectively, were generated using custom R script.

### ATAC-Seq

10^5^ cells were lysed with ATAC buffer (Tris 10 mM, pH 7.4, 10 mM NaCl, 3 mM MgCl2, NP-40 0.1%) and nuclei were collected for tagmentation at 37 °C for 30 minutes. The reaction was stopped with 0.2% SDS and DNA was purified using Qiaquick PCR Purification Kit (Qiagen) and eluted in 10 μl water. Eluted DNA was amplified using NEBNext Ultra II PCR Master Mix (NEB) and purified using AMPure XP beads. Samples were pooled for multiplexing and sequenced using paired-end sequencing on the Illumina NextSeq 500.

#### ATAC-seq analysis

ATAC-seq datasets for H3.3 KO, HIRA KO, ATRX KO, DAXX KO and corresponding WT cells were obtained from GEO (GSE151013) [80]. FastQ reads were trimmed and adapters removed using Trimgalore and Cutadapt. Quality of reads was assessed using FastQC. ATAC-seq reads were aligned to the mouse reference genome (mm10) using Bowtie2 [81]. Optical duplicates were removed using Picard and reads which mapped to the mitochondrial were filtered out. Peak calling was performed using MACS2 software [79] with a *p* value cutoff of 0.01. Peaks were further filtered to remove blacklisted regions using BEDTools [77]. A merged peak file containing all filtered ATAC peaks in every sample was used for downstream TOBIAS analysis. MACS2 was further used to generate bedgraph files which were normalized to signal per million reads. UCSC-userApps was used to convert bedgraphs to bigWigs for visualization in IGV [78]. Heatmaps and average profiles were generated from bigWig files using deepTools [82]. Differentially accessible peaks were identified using the DiffBind package [83].

#### Motif analysis

Active genes and expressed transcription factors were identified from RNA-seq [25] using a cut-off of >20 baseMean from DESeq2. Frequency matrices for each TF motif were downloaded from JASPAR database [84]. The TSS for active genes was downloaded as a bedfile from UCSC Table Browser and further expanded to 3 Kb on either side. ATAC-seq peaks that intersected TSS beds were identified as promoter-proximal. Motifs within promoter-proximal ATAC peaks were identified using FIMO [85].

#### NucleoATAC analysis

For NucleoATAC analysis, replicate bam files were merged and normalized to the same sequencing depth using Samtools. Nucleosome positioning profiles were generated from merged bam files around active promoter regions (TSS ± 1 kb) with default setting of nucleoATAC. Genome browser tracks were generated converting nucleoatac_signal.smooth.bedgraph to bigwig format using bedGraphToBigWig.

#### Tobias analysis

For TOBIAS analysis, replicate bam files were merged. TOBIAS ATACorrect and ScoreBigWig were used to generate scored bigWig files for each merged sample across the merged ATAC peaks bedfile. BINDetect was then used to generate pairwise differential binding scores between samples for each expressed JASPAR motif. For analysis of differential binding scores specifically in promoters, BINDetect was restricted using option --output-peaks to active promoter regions (TSS ± 3 kb).

### Global run-on sequencing (GRO-seq)

ESCs were lysed in Hypotonic Lysis Buffer [10 mM Tris pH 7.4, 0.5% NP-40, 10% glycerol, 3 mM CaCl2, 2 mM MgCl2, 1 mM DTT, 1x protease inhibitor cocktail (Roche), and SUPERase-In (Thermo Fisher)]. Nuclei were collected by centrifugation, washed once with 1 mL Lysis Buffer and resuspended in 500 µL of Freezing Buffer (50 mM Tris pH 8.3, 40% glycerol, 5 mM MgCl2, 0.1 mM EDTA, and 4 units/mL of SUPERase-In per mL) and stored at −80 °C.

For nuclear run-on, 5×10^6^ nuclei in 100 µl Freezing Buffer were mixed with an equal volume of 2X Run-on mastermix [(10 mM Tris pH 8.0, 2.5 mM MgCl_2_, 0.5 mM DTT, 150 mM KCl, 0.25 mM rATP, 0.25 mM rGTP, 1 µM rCTP, 0.25 mM bromo-UTP, 1% sarkosyl and 0.1 U/µl SUPERaseIn (Thermo Fisher)] and incubated at 30 °C for 5 minutes. The reaction was stopped by treatment with DNaseI and Proteinase K. NaCl was added to 225 mM, and the reaction was extracted twice with acid phenol:chloroform and once with chloroform. Following precipitation, RNA was hydrolyzed with 1N NaOH for 15 minutes on ice, and treated with DNaseI and PNK. Fragmented RNA was bound to anti-BrdU beads in binding buffer [37.5 mM NaCl, 1 mm EDTA, 0.05% Tween, 0.25× saline-sodium-phosphate-EDTA buffer (SSPE)] for 1 h at room temperature. Beads were washed once with binding buffer, low salt wash buffer (0.2X SSPE, 1 mM EDTA, 0.05% Tween) and high salt wash buffer (0.25X SSPE, 1 mM EDTA, 0.05% Tween, 137.5 mM NaCl), followed by two washes in TET buffer (TE + 0.05 mM Tween). RNA was eluted 4x with elution buffer (50 mM Tris pH 7.5, 150 mM NaCl, 1 mM EDTA, 20 mM DTT, 0.1% SDS) and extracted with acid phenol:chloroform. The resulting RNAs were reverse transcribed, size-selected and amplified. The quality of the libraries was assessed using a D1000 ScreenTape on a 2200 TapeStation (Agilent) and quantified using a Qubit dsDNA HS Assay Kit (Thermo Fisher). Libraries with unique adaptor barcodes were multiplexed and sequenced on an Illumina NextSeq 500.

#### GRO-seq analysis

Quality control for the GRO-seq data was performed using the FastQC tool. GRO-seq reads were trimmed to remove adapter contamination and poly(A) tails using the default parameters of Cutadapt software (Martin 2011). Reads >32 bp long were retained for alignment to the mouse reference genome. Transcript calling was performed using groHMM.

### RNA-seq analysis

RNA-seq datasets used for this study were obtained from the GEO (GSE114549). Quality of raw RNA-seq reads was assessed using the FastQC tool. Reads were aligned to the mouse reference genome (mm10) with STAR [86]. After normalization, the reads were converted into bigWig files using BEDTools [77] for visualization in Integrative Genomics Viewer or the UCSC genome browser. Count matrices were generated using the featureCounts tool [87] and differential expression analysis was performed using DESeq2 (version 1.14.1) with FDR cutoff p<0.05.

MA Plots. For each comparison, the mean of normalized counts and the log_2_ fold change of read counts in each gene were determined using DESeq2. These values were plotted on the x axis and y axis respectively for all transcripts detected. To visualize changes in expression of genes thought to play a role in pluripotency, mouse homologs of genes included in the core ESC-like gene module [56] or PluriNet [57] were highlighted.

### Western Blotting

ESCs were lysed with micrococcal nuclease (Worthington) in 50 mM Tris pH 7.5, 1 mM CaCl_2_, 0.2% Triton X-100 and 5 mM sodium butyrate. Proteins from whole cell lysate were separated in Laemmli buffer by SDS-PAGE and transferred to PVDF membranes (Millipore). Membranes were blocked in 5% milk in Tris-buffered saline with 0.1% Tween-20 (TBST) and incubated with primary antibodies overnight at 4 °C or for 2 hours at room temperature. Membranes were washed with TBST, incubated with HRP-conjugated secondary antibodies for 1 h, incubated with HRP substrate (Fisher) and imaged using a ChemiDoc MP Imaging System (BioRad).

## Declarations

### Ethics Approval and Consent to Participate

Not applicable.

### Consent for Publication

Not applicable.

### Availability of Data and Materials

**Code Availability.** Code to generate figures is available at https://github.com/utsw-medical-center-banaszynski-lab/Tafessu-et-al-2021

**Data Availability.** Datasets are deposited in the NCBI Gene Expression Omnibus using the following accession numbers: SuperSeries GSExxx, ATAC-seq GSExxx, ChIP-seq GSExxx, and GRO-seq GSExxx. Other datasets used for this study are available under GSE114551 [25] and GSE151058 [80].

## Authors’ Contributions

A.T., R.O., and S.M. contributed equally to this work. A.T., R.O., S.M., and L.B. conceived and developed the project. A.T., S.M., and L.A.B. designed the experiments and oversaw their execution, with assistance from A.L.D. and P.S. A.T. and S.M. performed the ATAC-seq, ChIP-seq, and GRO-seq experiments. A.T., R.O., and S.M. analyzed the data and performed integrative analysis of genomic data sets. A.T. prepared the initial draft of the text, which was edited by R.O., finalized by L.A.B., and approved by all co-authors. L.A.B secured funding for the project and provided intellectual support for all aspects of the work.

## Acknowledgements

We thank members of the Banaszynski lab for feedback on this project. We thank UTSW BioHPC for computational infrastructure and UTSW McDermott Center for providing next-generation sequencing services.

---

**Supplemental Figure 1. Related to Figure 1.**
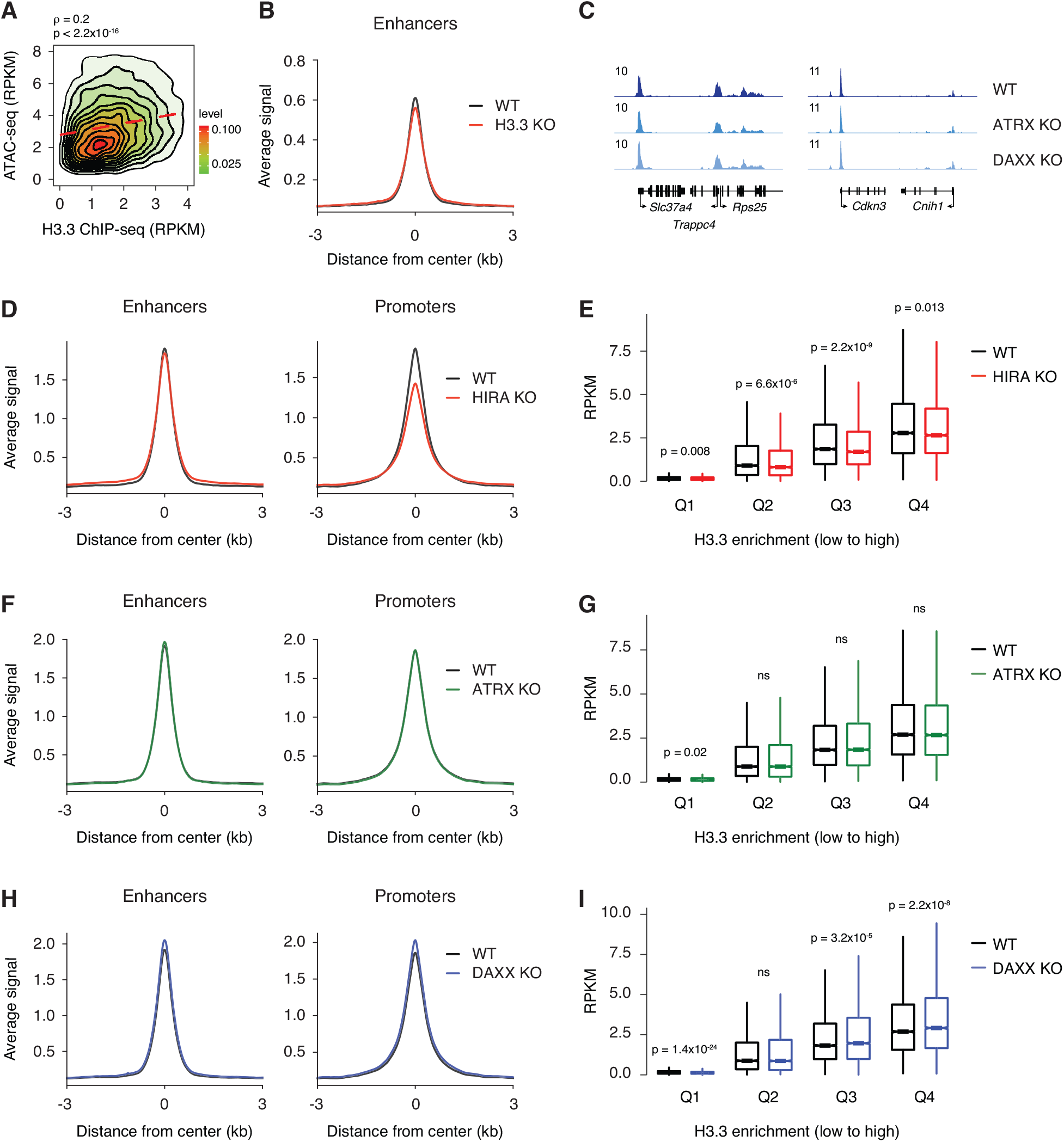
Loss of H3.3 deposition reduces chromatin accessibility at promoters. **A** Correlation plot between ATAC-seq and H3.3 ChIP-seq at enhancers in ESCs. **B** ATAC-seq average profiles at enhancers in WT and H3.3 KO ESCs. **C** Genome browser representations of ATAC-seq in WT, ATRX KO, and DAXX KO ESCs. The y-axis represents read density in reads per kilobase per million mapped reads (RPKM). **D, F, H** ATAC-seq average profiles at promoters (left) and enhancers (right) in WT and **(D)** HIRA KO, **(F)** ATRX KO, or **(H)** DAXX KO ESCs. **E, G, I** Boxplot showing ATAC-seq signal at promoters binned by H3.3 enrichment in WT and **(E)** HIRA KO, **(G)** ATRX KO, or **(I)** DAXX KO ESCs. For all boxplots, the bottom and top of the boxes correspond to the 25th and 75th percentiles, and the internal band is the 50th percentile (median). The plot whiskers correspond to 1.5x interquartile range and outliers are excluded. P-values determined by Wilcoxon rank sum two-side test.

**Supplemental Figure 2. Related to Figure 1.**
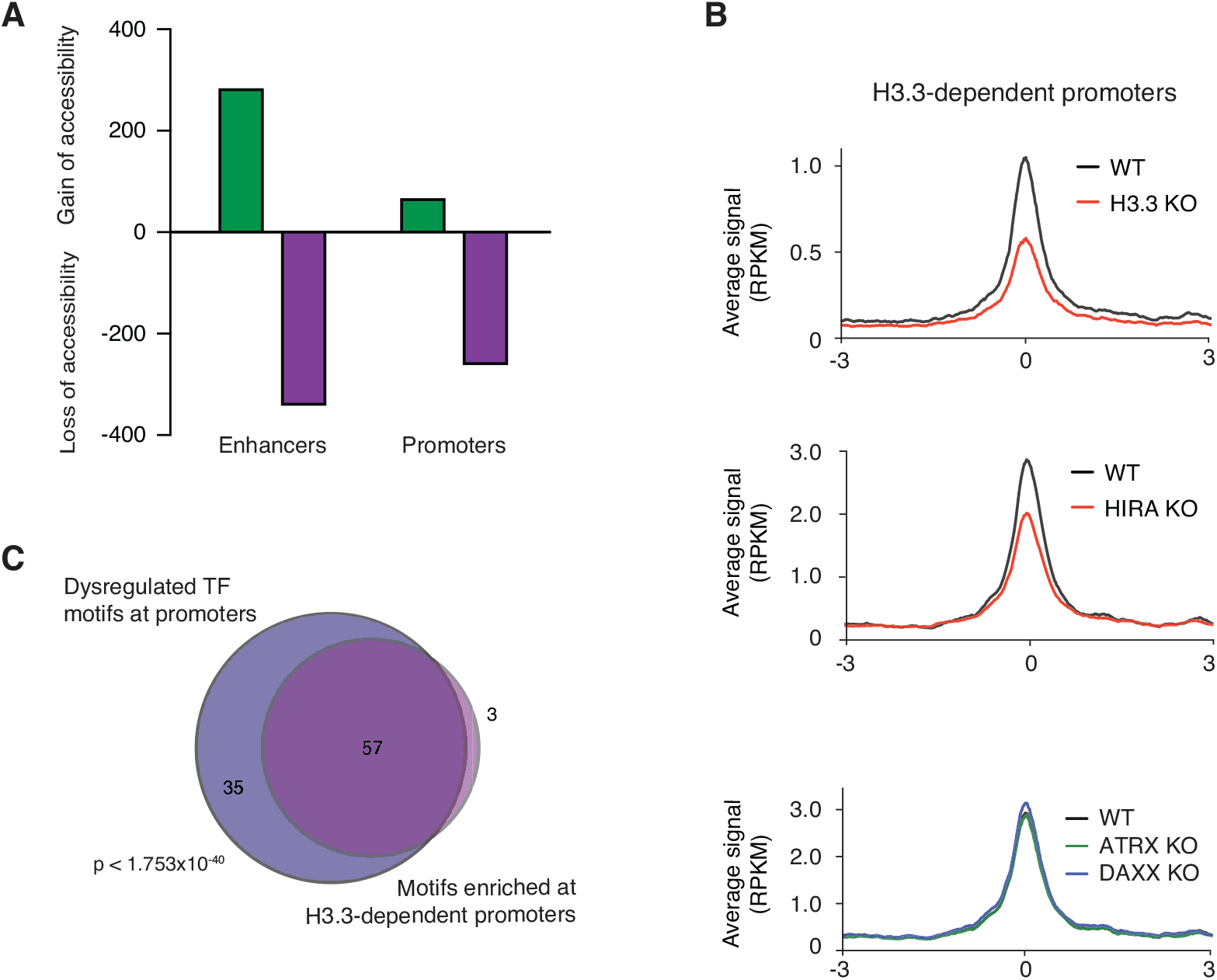
Promoter chromatin accessibility is facilitated by HIRA-dependent H3.3 deposition. **A** Representation of the number of differentially accessible enhancers and promoters in H3.3 KO compared to WT ESCs, determined using DiffBind. **B** ATAC-seq average profiles at H3.3-dependent promoters (i.e., promoters that lose accessibility in H3.3 KO compared to WT ESCs) in WT and H3.3 KO (top), HIRA KO (middle), and ATRX or DAXX KO (bottom) ESCs. **C** Venn diagram representing overlap between the top dysregulated motifs at promoters in H3.3 KO ESCs classified based on Manhattan scores in the top 10% across all comparisons (i.e., WT vs H3.3 KO, HIRA KO, ATRX KO, or DAXX KO) and TF motifs enriched at H3.3-dependent promoters.

**Supplemental Figure 3. Related to Figure 1.**
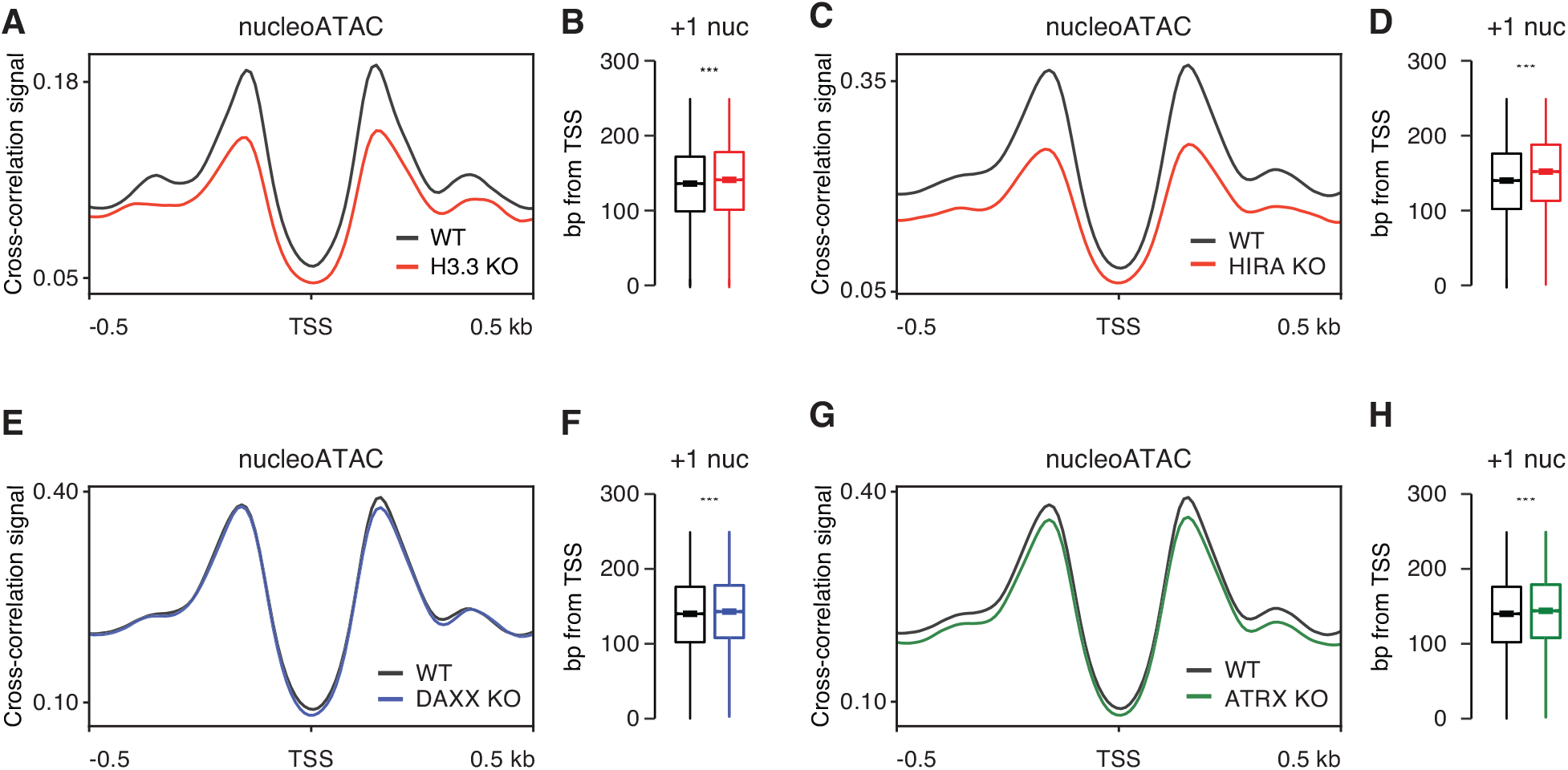
Loss of H3.3 alters promoter architecture. **A, C, E, G** Positive NucleoATAC cross-correlation signal at the TSS of active genes in WT and **(A)** H3.3 KO, **(C)** HIRA KO, **(E)** ATRX KO, and **(G)** DAXX KO ESCs. **B, D, F, H** Boxplot representing distribution of the +1 nucleosome from the TSS in WT and **(B)** H3.3 KO, **(D)** HIRA KO, **(F)** ATRX KO, and **(H)** DAXX KO ESCs. The bottom and top of the boxes correspond to the 25th and 75th percentiles, and the internal band is the 50th percentile (median). The plot whiskers correspond to 1.5x interquartile range and outliers are excluded. P-values determined by Wilcoxon rank sum two-side test.

**Supplemental Figure 4. Related to Figure 2.**
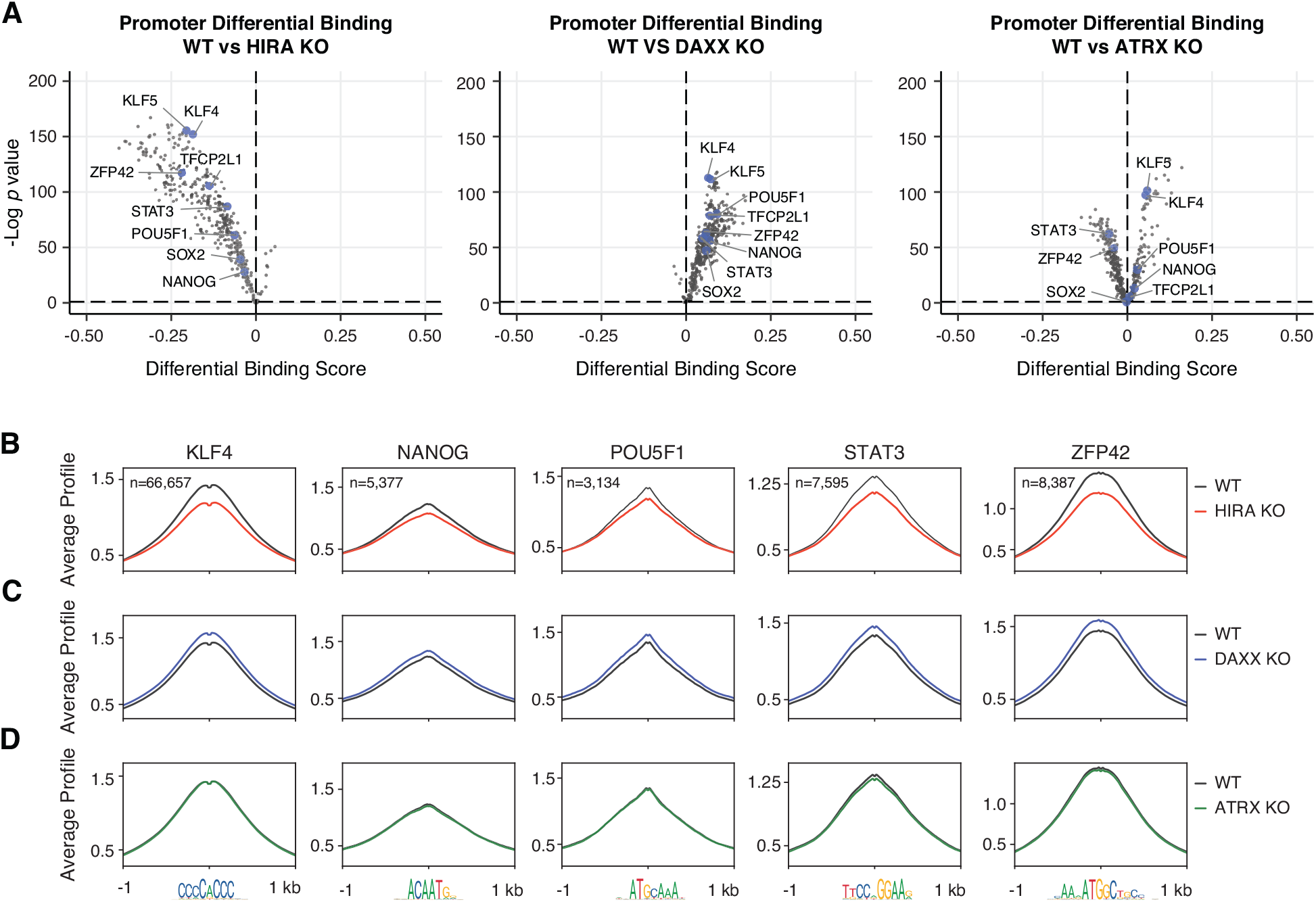
Loss of HIRA phenocopies promoter dysregulation. **A** Pairwise comparison of TF activity at promoters between WT and HIRA KO (left), DAXX KO (center), and ATRX KO (right) ESCs. Each TF is represented by a single circle (n = 395). TF motifs enriched in WT ESCs have negative differential binding scores and TF motifs enriched in chaperone KO ESCs have positive differential binding scores. Motifs for a subset of pluripotency-associated TFs are highlighted in blue. **B-D** ATAC-seq average profiles at representative TF motifs at promoters in WT and **(B)** HIRA KO, **(C)** DAXX KO, and **(D)** ATRX KO ESCs. Data are centered on the motif and the number of motifs profiled are indicated.

**Supplemental Figure 5. Related to Figure 2.**
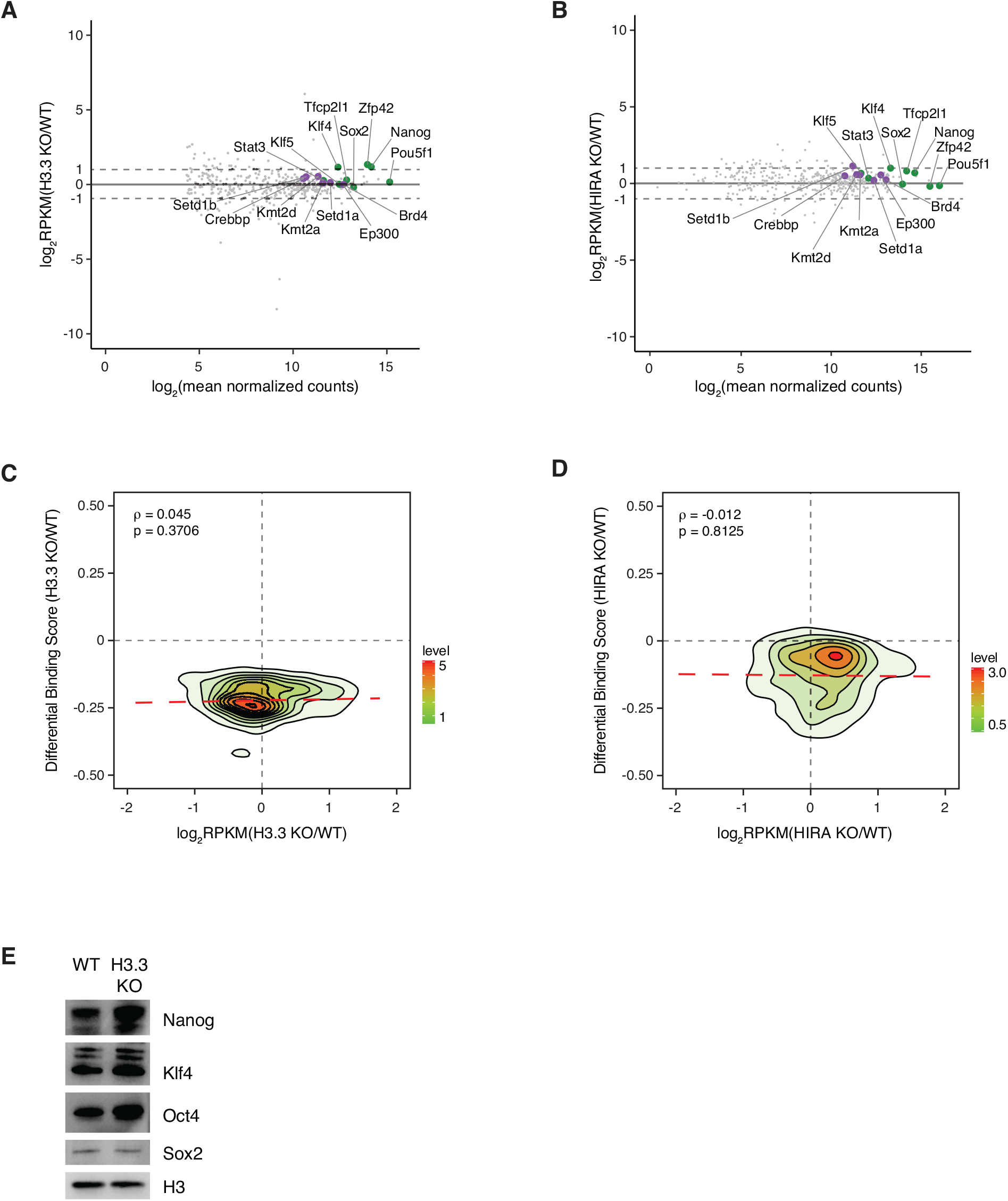
Changes in promoter architecture are not correlated with transcriptional changes in associated proteins. **A, B** MA plot representing all TFs represented in motif analysis and select chromatin-associated proteins. Mean expression across compared samples is represented on the x-axis and differential expression between WT and **(A)** H3.3 KO or **(B)** HIRA KO ESCs is represented on the y-axis. Highlighted TFs are labeled green and all chromatin-associated proteins are labeled purple. **C, D** Correlation plot between differential expression in WT and **(C)** H3.3 KO or **(D)** HIRA KO ESCs and differential TF binding score in WT and H3.3 KO ESCs. **E** Immunoblot of whole cell lysates from WT and H3.3 KO ESCs showing expression levels of NANOG, KLF4, OCT4 and SOX2. Total histone H3 was used as a loading control.

**Supplemental Figure 6. Related to Figure 3.**
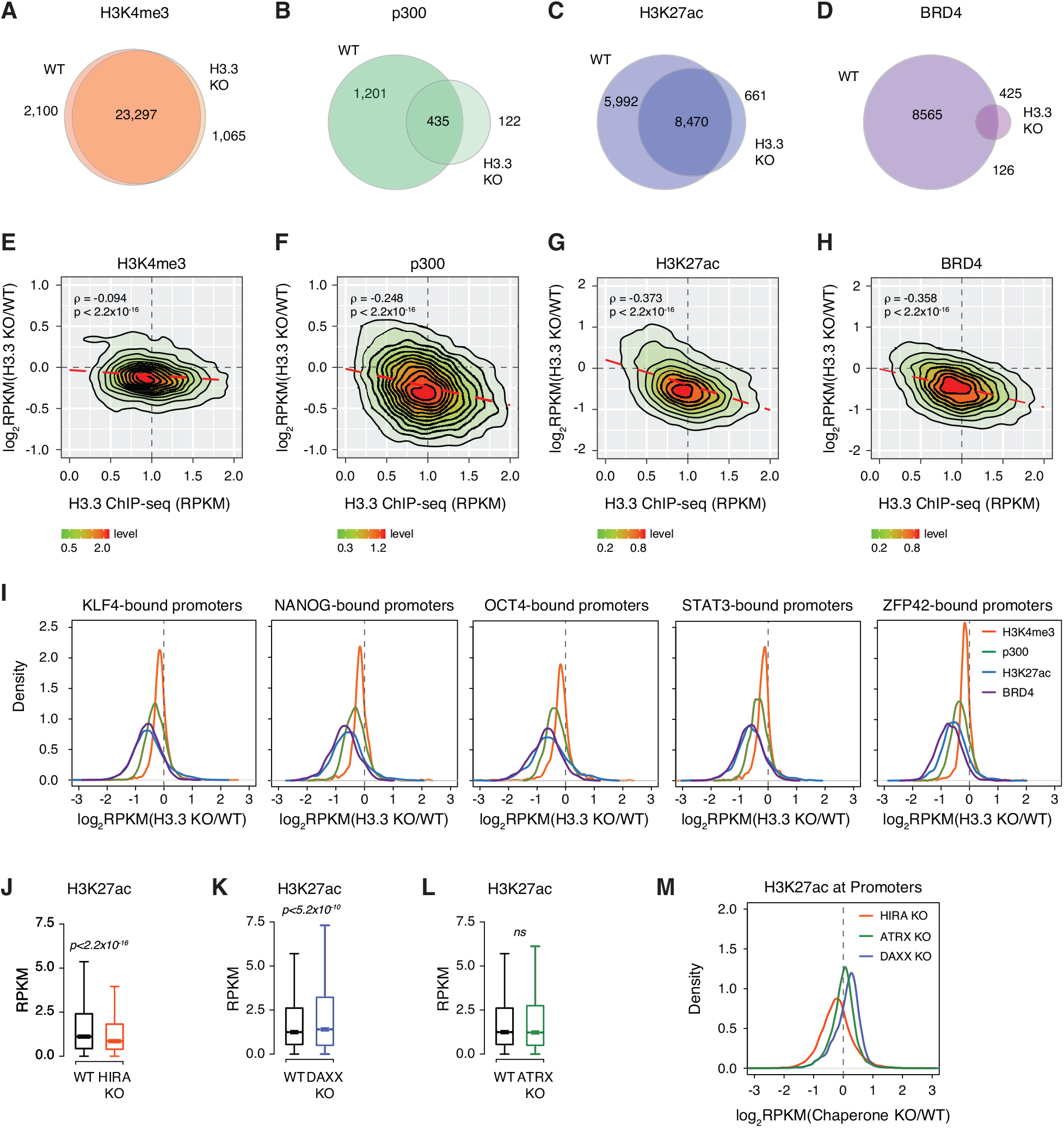
Chromatin landscape is dysregulated with H3.3 loss. **A-D** Venn diagram showing overlap between promoters enriched with **(A)** H3K4me3, **(B)** p300, **(C)** H3K27ac, and **(D)** BRD4 in WT and H3.3 KO ESCs. **E-H** Correlation plot between differential **(E)** H3K4me3, **(F)** p300, **(G)** H3K27ac, and **(H)** BRD4 enrichment in H3.3 KO compared to WT ESCs and H3.3 enrichment at promoters in WT ESCs. **I** Ratio (log2) of H3K4me3, p300, H3K27ac, and BRD4 enrichment at promoters bound by the indicated TF in WT and H3.3 KO ESCs. x axis values <0 indicate reduced enrichment in the absence of H3.3. **J-L** Boxplots showing H3K27ac enrichment at promoters in WT and **(J)** HIRA KO, **(K)** DAXX KO, and **(L)** ATRX KO ESCS (n = 12,903). The bottom and top of the boxes correspond to the 25th and 75th percentiles, and the internal band is the 50th percentile (median). The plot whiskers correspond to 1.5x interquartile range and outliers are excluded. P-values determined by Wilcoxon rank sum two-side test. **M** Ratio (log2) of H3K27ac enrichment at promoters in WT, HIRA KO, ATRX KO, and DAXX KO ESCs. x axis values <0 indicate reduced enrichment in the absence of chaperone.\

**Supplemental Figure 7. Related to Figure 5.**
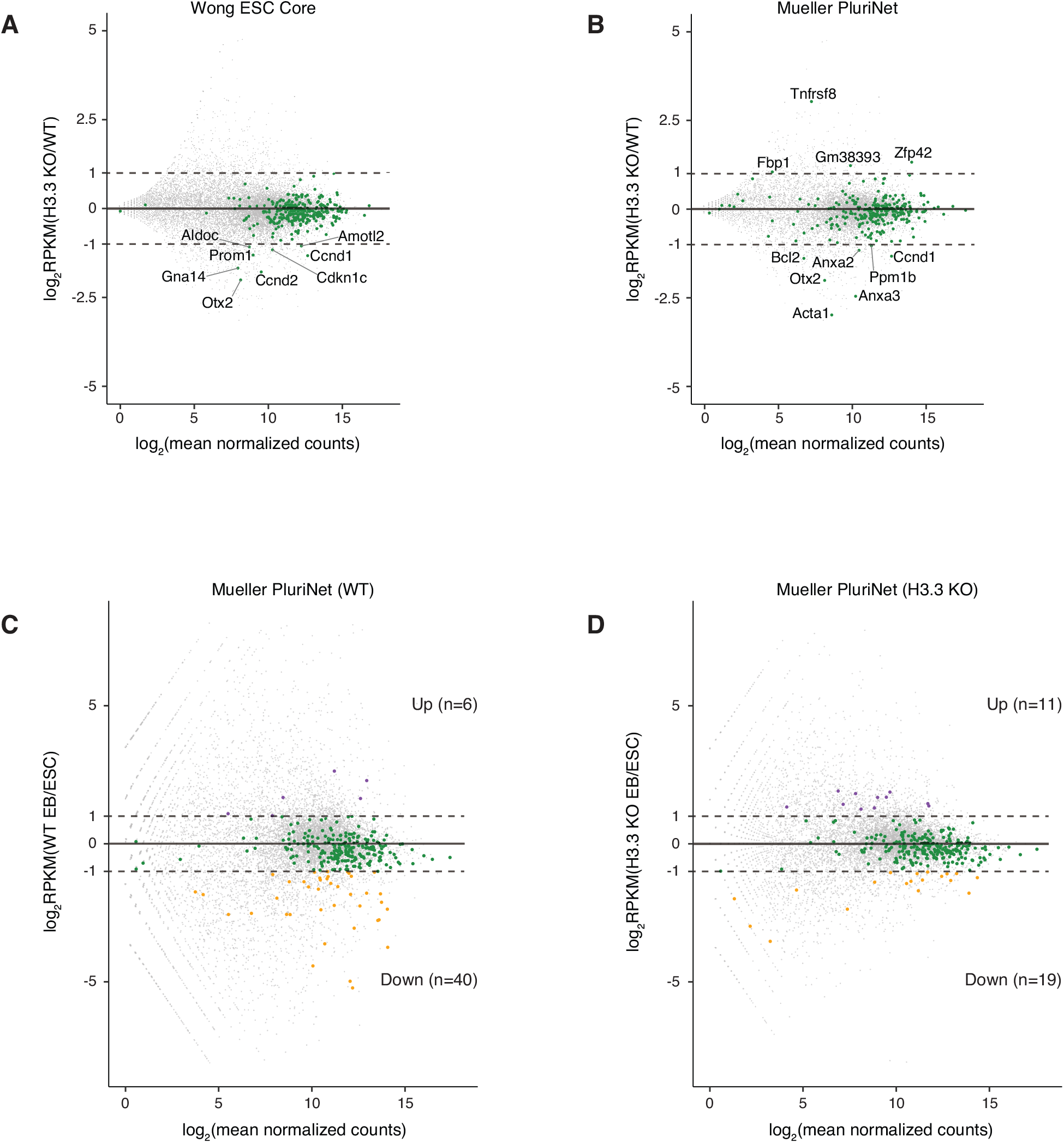
Transcriptional changes associated with loss of H3.3 in ESCs and during differentiation. **A-B** MA plot of gene expression in WT and H3.3 KO ESCs. Members of **(A)** the core ESC-like gene module or **(B)** PluriNet are shown in green. Mean expression across compared samples is represented on the x-axis and differential expression between WT and H3.3 KO ESCs is represented on the y-axis. **C-D** MA plot of gene expression in **(C)** WT and **(D)** H3.3 KO ESCs and EBs. Mean expression across compared samples is represented on the x-axis and differential expression between ESCs and EBs is represented on the y-axis. Members of the PluriNet gene set are shown in green (<2-fold change), purple (>2-fold increase in EBs) or orange (>-2-fold decrease in EBs).

